# Endemism shapes viral ecology and evolution in globally distributed hydrothermal vent ecosystems

**DOI:** 10.1101/2024.07.12.603268

**Authors:** Marguerite V. Langwig, Faith Koester, Cody Martin, Zhichao Zhou, Samantha B. Joye, Anna-Louise Reysenbach, Karthik Anantharaman

## Abstract

Viruses are ubiquitous in deep-sea hydrothermal environments, where they exert a major influence on microbial communities and biogeochemistry. Yet, viral ecology and evolution remain understudied in these environments. Here, we identified 49,962 viruses from 52 globally distributed hydrothermal vent samples (10 plumes, 40 deposits, and 2 diffuse flow), and reconstructed 5,708 viral metagenome-assembled genomes (vMAGs), the majority of which were bacteriophages. Hydrothermal viruses were largely endemic. However, some viruses were shared between geographically separated vents, predominantly between the Lau Basin and Brothers Volcano in the Pacific Ocean. Geographically distant viruses often shared proteins related to core functions such as structural proteins, and rarely, proteins of auxiliary functions. Common microbial hosts of viruses included members of Campylobacterota, Alpha-, and Gammaproteobacteria in deposits, and Gammaproteobacteria in plumes. Campylobacterota- and Gammaproteobacteria-infecting viruses reflected variations in hydrothermal chemistry and functional redundancy in their predicted microbial hosts, suggesting that hydrothermal geology is a driver of viral ecology and coevolution of viruses and hosts. Our study indicates that viral ecology and evolution in globally distributed hydrothermal vents is shaped by endemism, and thus may have increased susceptibility to the negative impacts of deep-sea mining and anthropogenic change in ocean ecosystems.

## Introduction

An estimated 10^30^ viruses in the world’s oceans are predicted to lyse and kill 20% of microbial biomass daily^1^. In marine systems, global sampling efforts have resulted in the recovery of viruses at a broad scale, enabling investigations of their ecology, evolution, and biogeography^2–4^. These studies have revealed that viruses in the epipelagic ocean are passively transported through ocean currents^5^, distinct viral groups exist across five oceanic ecological zones^4^, and there are broad differences in the protein content of viruses with depth^6^. Though these studies provide valuable insights on viruses globally, they have mainly focused on comparative analyses of photic and aphotic viruses. Investigations of viruses in the deep ocean are lacking.

In deep-sea hydrothermal vents, viruses are an important source of predation on chemolithoautotrophic microorganisms that serve as the base of the food web in the absence of sunlight^7^. As a result of these infections, viruses have the potential to change microbial community composition and population sizes. Hydrothermal vent viruses are also capable of “reprogramming” host metabolism using auxiliary metabolic genes (AMGs). For example, vent viruses were the first viruses discovered to encode reverse dissimilatory sulfite reductase (rdsr), which can be used to manipulate sulfur-based chemolithotrophy in the dark ocean^8^. To date, few studies have conducted comparative analyses of hydrothermal vent viruses globally, and those completed provide evidence that viruses are endemic to vent sites and habitat types at the genus level, and that they infect ecologically important, abundant taxa such as Gammaproteobacteria and Campylobacterota^9^. Other studies have suggested vent viruses are predominantly lysogenic^10^, have limited dispersal, and narrow host ranges^11^.

Despite these advances, the factors controlling hydrothermal vent virus biogeography, ecology, and evolution remain poorly constrained. The biogeography of viruses is thought to be determined by a complex interplay between abiotic factors, virus traits (e.g., life cycle, virion size, burst size), and host traits (e.g., abundance, size, distribution)^12^. Abiotic factors such as vent geochemistry have been shown to dictate microbial community composition, where differences in the geochemical profiles of geographically close vent sites can result in distinct microbial communities^13,14^. Given their dependence on microbial hosts, viral biogeography is intimately linked to theories on microbial biogeography, however, these remain in their infancy^15,16^. Increasingly available hydrothermal vent metagenomic data presents opportunities to examine vent viral biogeography, ecology, and evolution at a global scale. This, coupled with advances in software for rapid, accurate comparison of metagenome-assembled genomes (MAGs)^17^ and the generation of viral MAGs (vMAGs)^18^, promise to enable more accurate representations of environmental viruses and allow finer resolution comparisons of their community structure and ecology.

In this study, we catalog and describe viruses, largely from two types of hydrothermal vent environments, on a global scale: high temperature hydrothermal deposits that host biofilms of thermophilic bacteria and archaea, and hydrothermal plumes hosting psychrophilic and mesophilic bacteria and archaea. These samples have previously been investigated for microbial diversity^13,14,19^, leaving their viral communities largely unexplored. Using 10 hydrothermal plume, 40 vent deposit, and 2 diffuse flow samples collected from seven distinct hydrothermal systems in the Pacific and Atlantic Oceans (Guaymas Basin, Mid-Cayman Rise, Mid-Atlantic Ridge, Axial Seamount, Brothers Volcano, East Pacific Rise, and the Lau Basin), we leveraged metagenomics and statistical analyses to reconstruct viral genomes and study viral communities through inter- and intra-vent comparative analyses.

## Results

We identified 63,826 viral scaffolds from 10 hydrothermal plumes, 40 hydrothermal vent deposits, and 2 diffuse flow samples (**Supplementary Table 1**). Following viral identification, we conducted viral genome binning to reconstruct 5,708 vMAGs. The vMAGs comprise 19,572 viral scaffolds (30.7% of scaffolds), leaving 44,254 unbinned viruses (69.3% of scaffolds). Thus, after viral binning, we recovered a total of 49,962 viruses from globally distributed hydrothermal vents (**Figure 1A**). Of these, 1,833 were characterized as medium-, high-quality, or complete (**Supplementary Figure 1**) and 20,305 viruses encoded one or more viral hallmark genes (**Supplementary Table 2**). Most of the hydrothermal vent viruses were classified as lytic rather than lysogenic (47,571 lytic versus 2,391 lysogenic) and this remains true when only examining viruses of medium-quality or better (1,505 lytic versus 328 lysogenic). In addition, 32,442 vent viruses had a genome size range of 1-5 kb, while the remaining had genome sizes of 6-561 kb (**Supplementary Figure 2**). Taxonomic predictions at the class-level showed that most viruses are double-stranded DNA viruses within the realm *Duplodnaviria* (39,056), class *Caudoviricetes* (**Supplementary Figure 3**). *Caudoviricetes* viruses were also the most abundant class of viruses in the dataset based on relative abundance (**Figure 1B, Supplementary Table 3**). Most viruses were reconstructed from deposit samples from Brother’s Volcano and the Lau Basin (35,094 viruses). These sites produced some of the largest assemblies of the datasets analyzed here (up to 1.2 Gb) and were the most intensively sampled compared to other sites.

**Figure 1.**
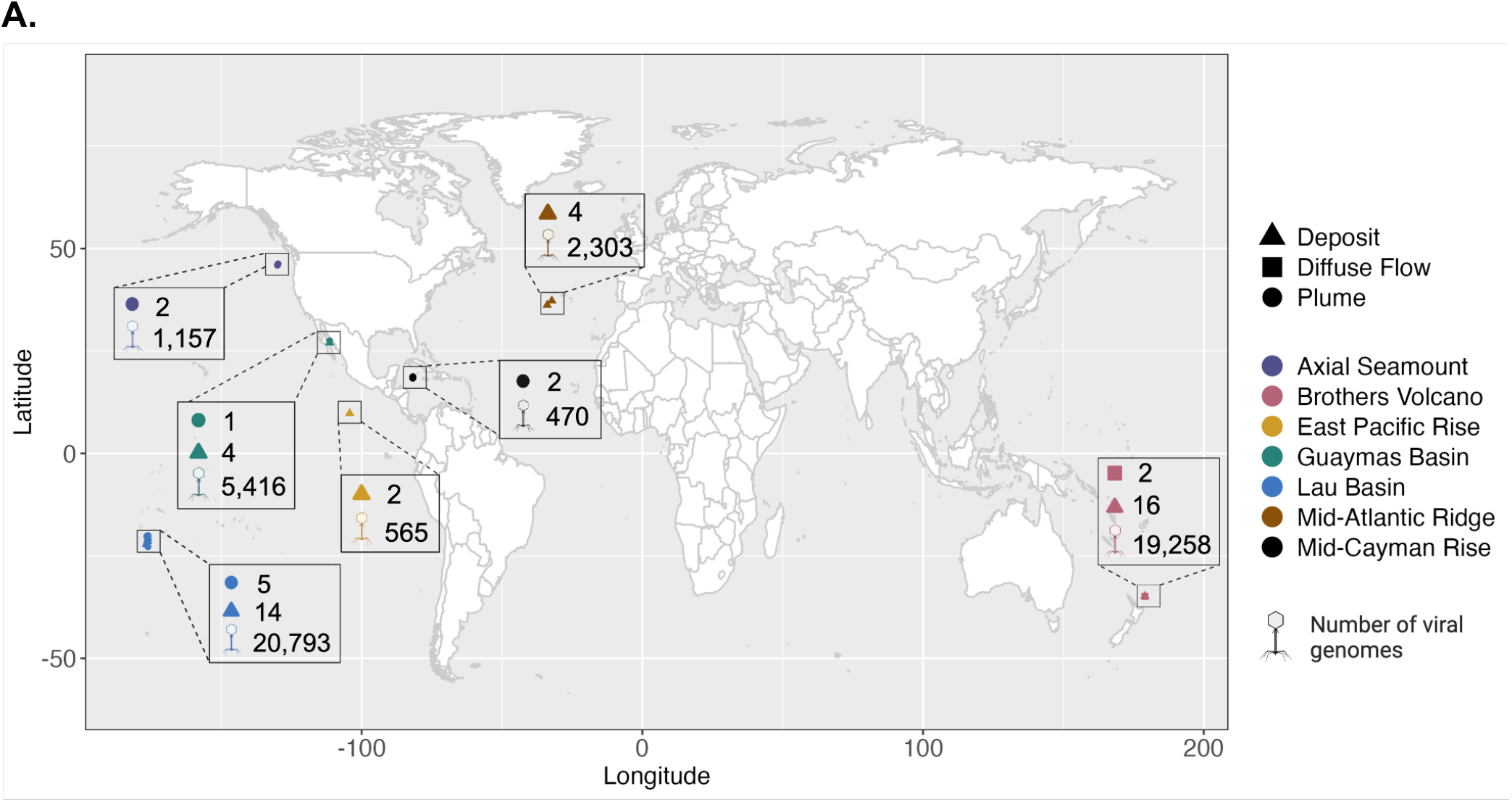
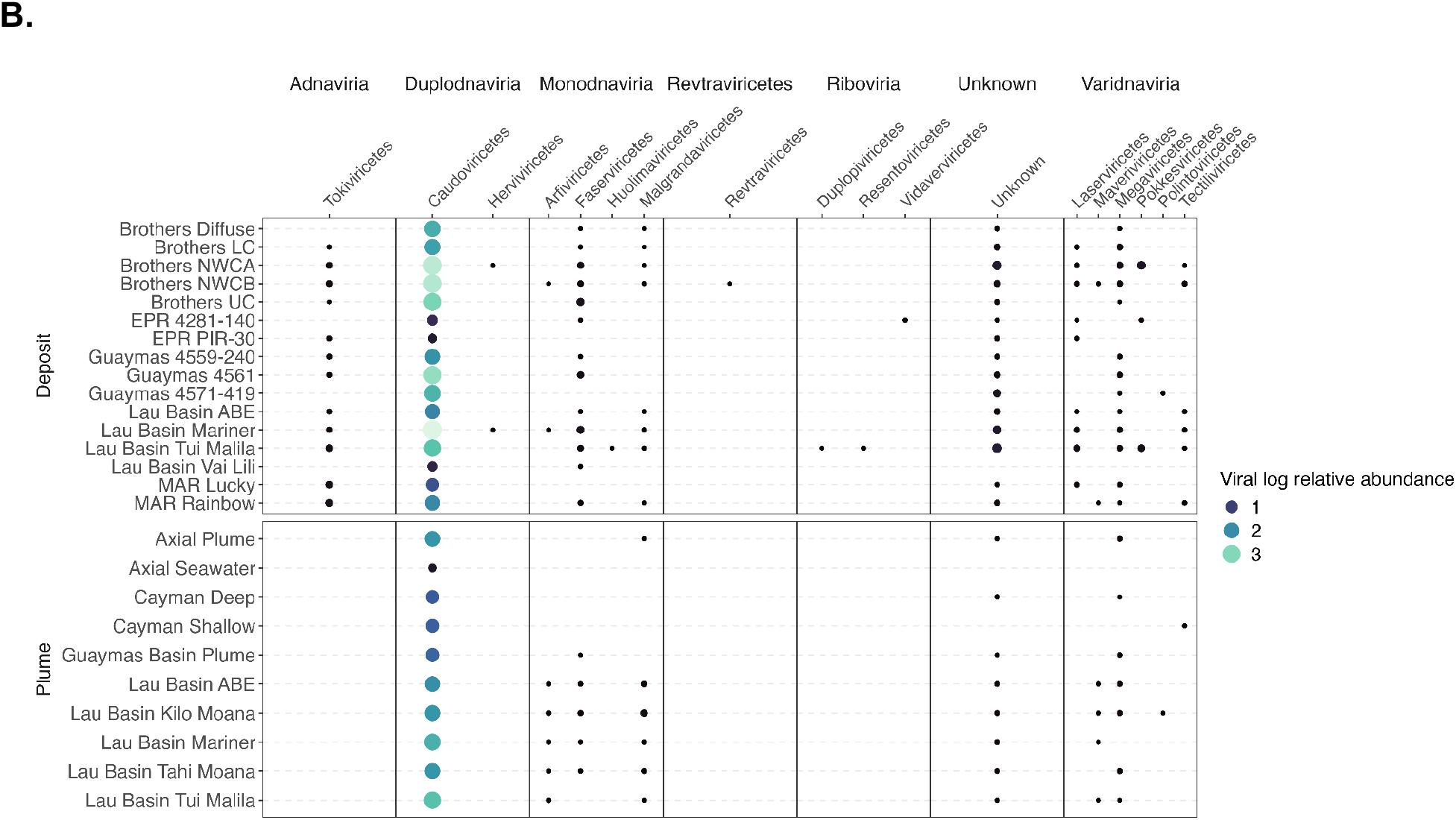
Geographic distribution, abundance, and taxonomy of viruses identified in globally distributed hydrothermal vents. **A**. A world map showing the number of viruses identified from different hydrothermal vents. Circles represent metagenomic samples reconstructed from hydrothermal plumes, triangles are metagenomic samples reconstructed from hydrothermal vent deposits, and squares are diffuse flow. Numbers next to the shapes represent the number of samples for that vent field (52 total). Virus icons show the number of viruses identified at a vent site (49,962 total). Colors represent the seven distinct hydrothermal vent fields that are shown in the legend. **B**. A bubble plot of log-transformed virus relative abundance, summed by viral class. Circles represent the log relative abundance, where larger relative abundance is represented by larger, yellow circles, and smaller relative abundance is represented by smaller, dark blue circles. Virus class names are shown on the top x axis and the horizontal names above them show virus realms (determined using geNomad). Site names are shown on the left y axis and the vertical names to the left of them show sample type.

### Geographically distant hydrothermal vents rarely share viruses

To understand how hydrothermal vent viruses are related, we conducted similarity analyses of viruses across and within hydrothermal vents. These analyses included clustering and network analysis based on average nucleotide identity (ANI), as well as read mapping of genomes. Clustering of viral genomes identified 866 non-singleton clusters containing 1,950 viruses, and most clusters (687/866) were composed of two viral genomes (**Supplementary Table 4**). Thus, most vent viruses were not included in ANI-based clusters, suggesting they have low relatedness at the nucleotide level. In addition, no clusters contained viruses from both hydrothermal plumes and hydrothermal deposits, indicating these habitats support distinct viral communities. Interestingly, some viruses fell within the same nucleotide clusters, yet were reconstructed from geographically distant sites (**Figure 2A**, red outlined ribbons). Specifically, 65 clusters contain 152 viruses from geographically distinct vent deposits, with an ANI ranging from 70-99%. When examining the clusters, we find that the predicted viral genome sizes, completeness, hosts, and lifestyles are largely aligned. Most of the clusters (51/65) contained viruses from the Lau Basin and Brothers Volcano, which are both located in the South Pacific Ocean. Viral genomes in these clusters shared significant overlap even across geographically separated vents in different ocean basins, such as between Mid-Atlantic Ridge in the Atlantic Ocean and Brothers Volcano in the Pacific Ocean, or between Axial Seamount and Guaymas Basin in the Pacific Ocean (**Supplementary Table 5**). Although viruses in the 65 geographically distinct clusters are predominantly predicted to be low-quality, we determined that the regions of overlap in these viruses are significant in length and/or annotation, and thus we find support for shared viral genomic regions between geographically separated vents (**See Supplementary Text**).

**Figure 2.**
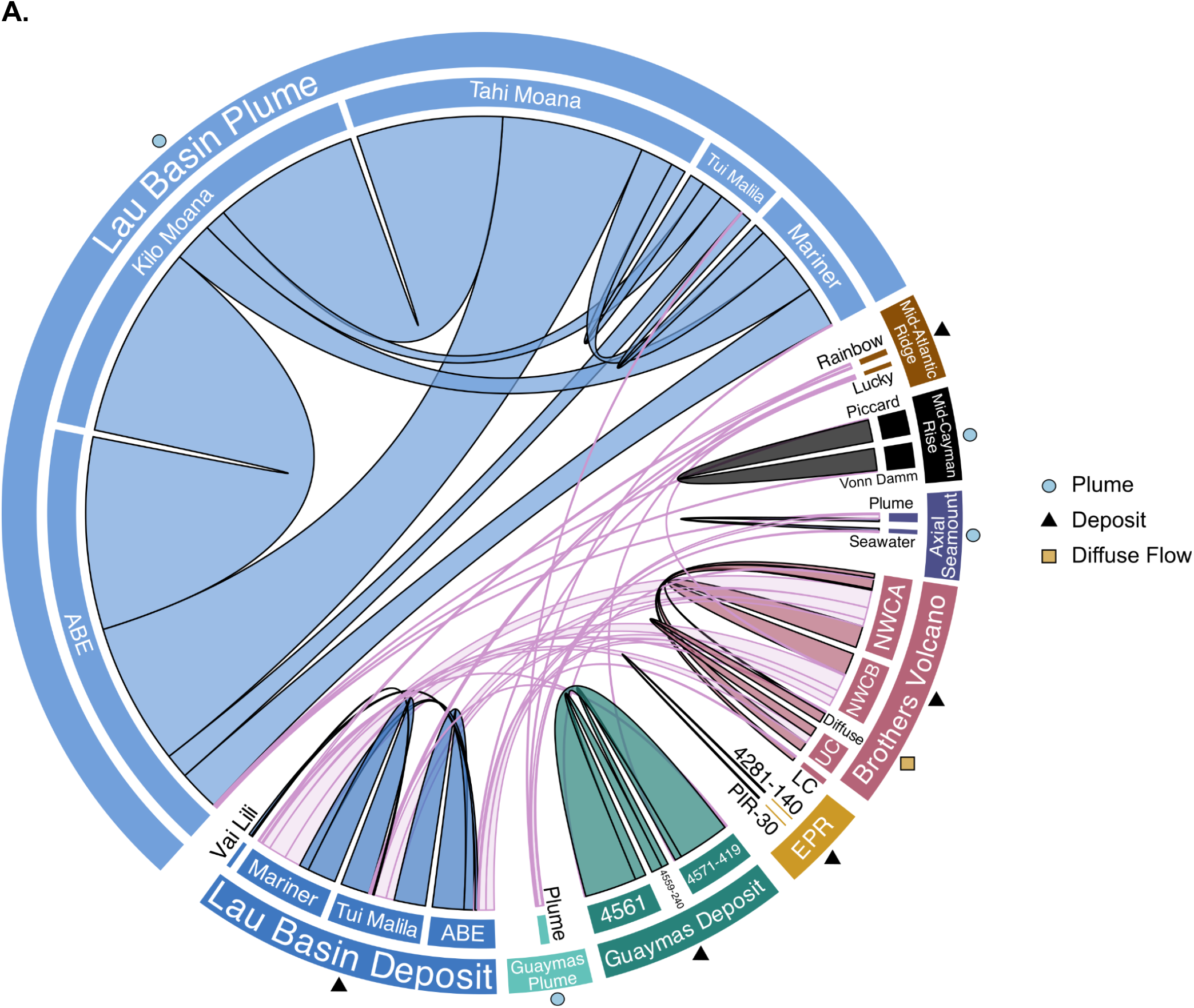
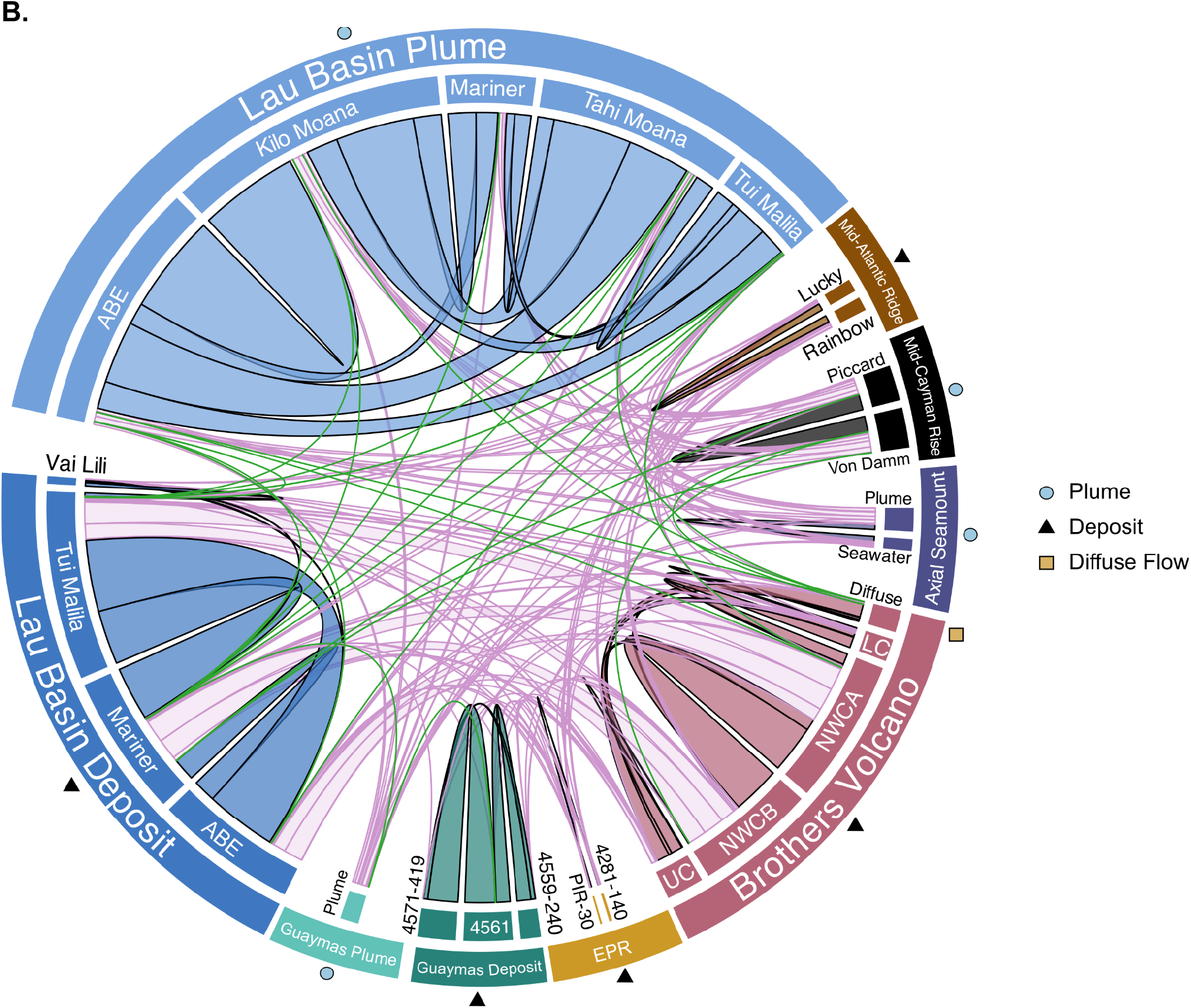
Biogeography of hydrothermal vent viruses. **A**. Viral relatedness based on average nucleotide identity of ≥3kb viruses and mcl clustering. Ribbons signify clusters that have viruses shared between hydrothermal vent sites (≥70% ANI). The width of the ribbon represents the number of clusters containing viruses from each site. Ribbons outlined in light purple show clusters with viruses from geographically distinct vent sites while ribbons outlined in black with the fill color of the site indicate clusters with intra vent viruses (viruses from the same vent field but distinct vent locations). On the outer ring, circles show plume samples, triangles signify deposit samples and squares show the diffuse sample. **B**. Viral relatedness based on read mapping between all reads and all viral genomes ≥3kb length and ≥70% coverage. The width of the ribbon represents the number of times reads from one site mapped to a virus from another site. Ribbons outlined in light purple highlight reads that mapped between a vent and a virus from geographically distinct sites and ribbons outlined in green highlight reads that mapped between vent plumes and deposits. Ribbons outlined in black with the fill color of the site indicate instances where reads from a vent site mapped to viruses from the same vent field (intra-vent read mapping). On the outer ring, circles show plume samples, triangles signify deposit samples and squares show the diffuse sample.

Some shared viruses may be missed when examining viruses identified from metagenomes, because viral sequences may only be present in the reads and not the assemblies. To address this issue, we used read mapping to identify viruses that were present in multiple samples. While nucleotide clustering indicated that no viruses were shared between hydrothermal vent plumes and deposits, read mapping-based detection identified 36 such viruses (**Figure 2B, Supplementary Table 6**). Most of these were identified from Lau Basin (26/36) deposits or plumes from the same or different sites in Lau Basin. This suggests that some viruses may be shared between vent deposits and plumes.

### Viruses are more similar within a hydrothermal field

In addition to understanding how viruses are related between geographically distant vents, we examined intra-vent (different sampling locations within the same vent field) viral relatedness, where 417 clusters contained 992 viruses from distinct sites within a vent field (**Figure 2A**, grey outlined ribbons, **Supplementary Table 4**). Just over half of the clusters (272/417) contained viruses that were related between hydrothermal plumes in Lau Basin, including Lau Basin ABE, Kilo Moana, Mariner, Tahi Moana, and Tui Malila. These plumes are at most separated by 1-2º latitude (Supplementary Table 1). In the Lau Basin, 163/272 plume viral clusters had an average intra-cluster ANI ≥85% and 90/272 had an average ANI ≥95%. This indicated that many of these Lau Basin plume viruses were related at the genus and species level, or for those with 100% identity and 100% completeness, were identical. Almost all the viruses in Lau Basin plume clusters were predicted to be lytic and were represented by Microviridae, Caudoviricetes, Inoviridae, Schitoviridae, Cressdnaviricota, and Demerecviridae.

Lau Basin vent deposit samples had the next highest number of viral clusters shared between different vent fields (94/417 clusters, **Supplementary Table 4**). Of the 94 clusters, 16 contained medium-quality or better viruses, and nearly all of these were *Caudoviricetes*. Guaymas Basin and Brothers Volcano vent deposits, respectively, also contained many viral clusters shared between different vent fields. In Guaymas Basin, most clusters are shared between the sites 4561 (380 and 384) and 4571-419 (38/47), which are geographically close but separated by a depth of 22 m. Similarly, in Brothers Volcano, many clusters were shared between Northwest Caldera Wall A (NWC-A) and Northwest Caldera Wall B and Upper Caldera Wall (NWC-B+UCW) (13/30), which are geographically close to each other (∼1,570 m distance), though they have distinct microbial communities^13^. Further, Mid-Cayman Rise, Axial Seamount, and the East Pacific Rise also had some intra-vent related viruses. To complement our clustering analyses, we also conducted read mapping-based detection which indicated that viruses were related between intra-vent sites. We found that viruses from Lau Basin plumes were highly related (especially Kilo Moana and Abe, Tahi Moana and Kilo Moana). We also identified many viruses as shared between Lau Basin deposits (Tui Malila, ABE, and Mariner), as well as between Brothers Volcano deposits (NWC-A and NWC-B), Guaymas Basin deposits (4571-419 and 4561), Mid-Atlantic Ridge deposits (Lucky and Rainbow), and Mid-Cayman Rise plumes (Von Damm and Piccard). Overall, read mapping-based detection reaffirmed viral relatedness between sites identified with ANI clustering, but also identified numerous connections that were not observed with the ANI clustering analysis. This was especially true for overlap between viruses in hydrothermal plumes and deposits, viruses from the Mid-Atlantic Ridge, and viruses between more geographically distant vent fields like Kilo Moana and Cayman Shallow plumes, since these patterns were uniquely observed with read mapping.

### Hydrothermal vents share viral protein families dominated by proteins of unknown function

To understand how hydrothermal vent viruses are related at the protein level, we clustered all 595,416 vent virus proteins. This produced 74,940 clusters of two or more virus proteins (**Supplementary Table 7**). Of these, 152 clusters contained proteins shared between vent deposits and plumes (773 proteins), and 23,351 clusters have proteins shared between geographically separated vents from different vent fields (84,223 proteins or 14.2% of the protein dataset; **Figure 3**). Of the 84,259 total proteins shared across distant sites or sample types, 40,645 were annotated (48.2%), largely as hypothetical or uncharacterized functions, viral terminases, DNA/RNA polymerases, capsid, baseplate, tail, and portal proteins. Of the 152 clusters shared between vent plumes and deposits, 89 contained proteins with annotations, and the top annotation categories included helix-turn-helix domains, domains of unknown function, phage tail tube protein, hypothetical proteins, nucleotide kinase, and essential recombination function protein. The largest cluster of proteins from geographically separated vents had 88 proteins from deposit samples, including Brother’s Volcano, the Lau Basin, and Mid-Atlantic Ridge. This cluster contained a protein of unknown function that is known to often be encoded in phage genomes (PF09343 or IPR011740). Several of the next largest protein clusters contained proteins from the same sites (Brother’s Volcano, Lau Basin, and Mid-Atlantic Ridge) that are of unknown function or an AAA domain (PF13479). Many clusters with proteins from distant sites were also functionally related to pyrimidine metabolism (e.g., dCTP deaminase, dTMP kinase, and dUTP diphosphatase), purine metabolism (phosphoribosylformylglycinamidine cyclo-ligase and phosphoribosylamine-glycine ligase (PurD), and nucleotide metabolism (ribonucleotide reductase). Several clusters contained phosphate starvation inducible proteins (PhoH), which have previously been identified as widespread in marine phages and proposed as a marker of marine phage diversity^20^. Interestingly, some protein clusters that were highly similar between viruses from geographically distant vents contained proteins that do not have core functions, including pyruvate formate lyases (PflA) and cobaltochelatase subunits CobS and CobT (**Supplementary Table 8**).

**Figure 3.**
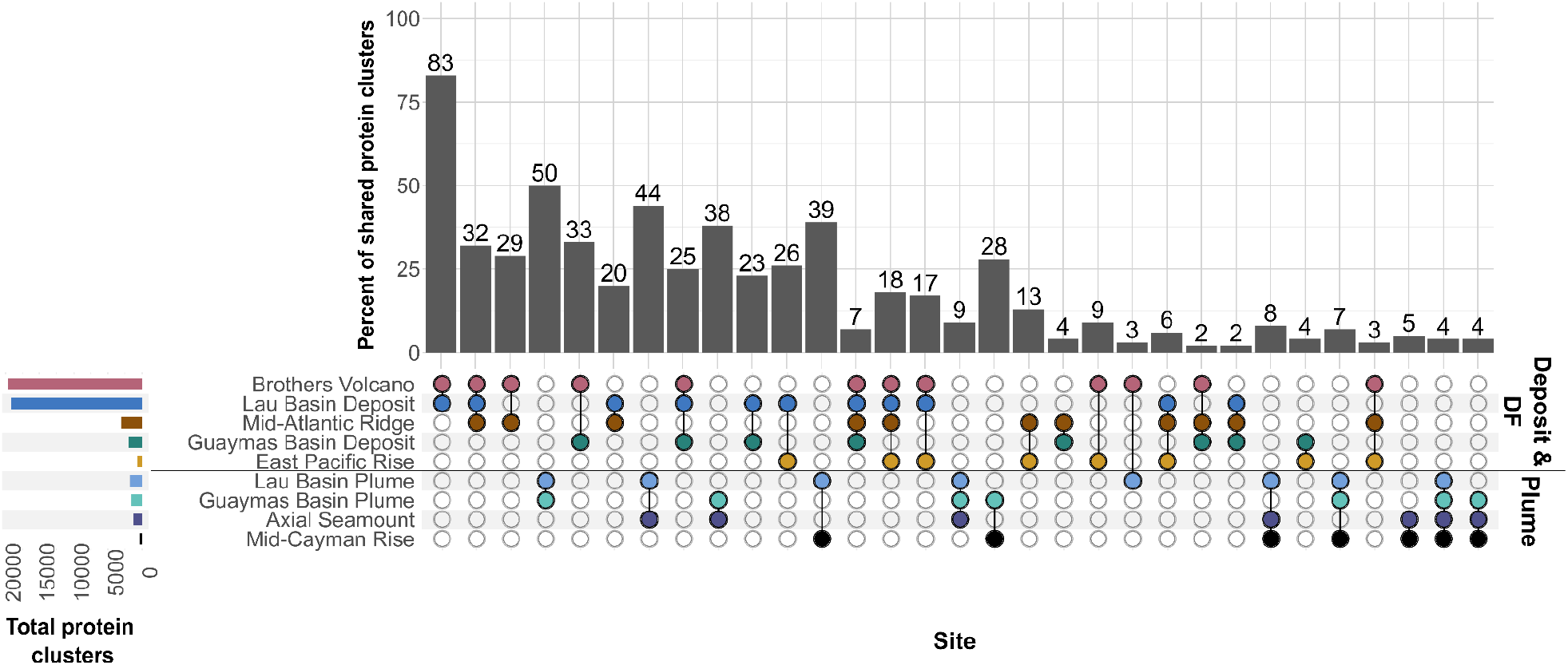
Shared viral proteins between geographically distant hydrothermal vents. The bar plot shows the percent of shared protein clusters, and the bottom matrix shows the identity of the sites with shared protein clusters (filled, colored circles). The percent of shared protein clusters was calculated as the number of shared protein clusters divided by the smallest number of total protein clusters for a group, multiplied by 100. The leftmost bar plot shows the total number of protein clusters per site. The black line through the matrix separates deposit and diffuse flow (DF) samples, shown as the first five sites, from plume samples, the bottom four sites. Sites with fewer than fifteen shared protein clusters were removed. All clusters are reported in Supplementary Table 7.

### Hydrothermal viruses encode auxiliary metabolic genes associated with redox processes and detoxification

The presence of auxiliary metabolic genes, or AMGs, in viruses may increase a virus’ potential geographic range^12^. Viruses that encode AMGs have an increased ability to boost energy levels for viral progeny production or reduce the viral latent period, and thus should be able to disperse more widely than viruses without AMGs. To investigate these dynamics, we searched for AMGs in all viruses (Supplementary Table 8). We also verified that AMGs were flanked by genes of viral or viral-like origin and were not present on the ends of genomic scaffolds. Important AMGs in hydrothermal vents were involved in sulfur metabolism, arsenic metabolism, nitrogen metabolism, and central carbon metabolism.

AMGs were rare in our dataset. According to DRAMv, 2,615 viruses encode one or more AMGs, or ∼5% of viruses recovered in this dataset. We identified a lytic *Caudoviricetes* virus reconstructed from Brother’s Volcano that encoded adenylylsulfate reductase (AprB, K00395), which catalyzes the reduction of adenylyl sulfate to sulfite in the dissimilatory sulfate reduction pathway, or the reverse reaction in dissimilatory sulfur oxidation. A lysogenic *Caudoviricetes* virus from Lau Basin Mariner encoded an arsenate reductase (ArsC, K00537), which functions in arsenate detoxification by reducing As(V) to an excretable form, arsenite or As(III)^21^. ArsC has been identified in soil viruses, where there is evidence that *arsC*-encoding viruses may contribute to metal resistance in their microbial host^22,23^. In addition to arsenate metabolism, we identified a virus encoding cytochrome bd ubiquinol oxidase subunit I and II (CydAB, K00425 and K00426), which acts as a terminal electron acceptor in the electron transport chain of microorganisms during respiration^24^. Although *cydAB* has not been described in other viruses, phage integration has been found to reprogram regulation of anaerobic respiration in *Escherichia coli*, and thus CydAB may be another mechanism by which viruses manipulate host respiration^25^. Finally, a *Caudoviricetes* virus from Lau Basin deposits was predicted to encode a nitric oxide reductase subunit B (NorB, K04561). NorB is the large subunit of nitric oxide reductase, which catalyzes the reduction of nitric oxide to nitrous oxide, the penultimate step of the denitrification pathway. NorB has previously been identified in viruses from an oxygen minimum zone in the Eastern Tropical South Pacific Ocean^26^.

### Viral biogeography is closely tied to the geographic distribution and abundance of their hosts

To explore microbial drivers of viral distribution, we predicted the microbial hosts and calculated the relative abundance of all viruses and their microbial hosts (**Supplementary Table 2, 3, and 9**). Of the 49,962 total viruses, 14% had a predicted host (7,001 viruses, **Supplementary Figure 4**). Virus infection range was largely narrow, where 6,387 viruses were predicted to infect one host, and the remaining 614 viruses were predicted to infect >1 host. Most host predictions were for deposit viruses (84.7%), which are predicted to infect a greater diversity and larger number of microbial phyla compared to viruses in plumes (**Supplementary Text**). Among plume viruses, most predicted hosts were members of the phyla Pseudomonadota (formerly Proteobacteria, 44.6%) and Bacteroidota (17.2%), while in deposits most were members of Campylobacterota (25%) and Pseudomonadota (21.2%, primarily Gamma- and Alphaproteobacteria). This aligned with relative abundance data, where the most abundant plume virus (Axial Plume, 1.3% relative abundance) infected a Proteobacteria in the class Gammaproteobacteria, genus *Thioglobus*, or SUP05. This sulfur-oxidizing bacterium is abundant in hydrothermal plumes globally^27,28^. In deposits and diffuse samples, the most abundant virus with a predicted host (Brothers Volcano site Diffuse, 0.9%) was predicted to infect a member of Campylobacterota in the family Sulfurimonadaceae (genus *CAITKP01*). Bacteria in this family and genus are the most abundant among all the deposit samples, and isolates in this family from hydrothermal vents are known to be chemolithoautotrophic sulfur oxidizers^14,29^.

### Hydrothermal geology and chemistry drive viral ecology and coevolution of viruses and hosts

In both deposits and plumes, microbial hosts from the phylum Pseudomonadota were largely associated with the class Gammaproteobacteria. Previously, using the same metagenomes in this study, functional redundancy was observed between members of Gammaproteobacteria and Campylobacterota, where these taxa shifted as dominant community members depending on the hydrothermal geology and chemistry. These microbial lineages have similar metabolic potential, and thus their dominance at one hydrothermal vent site or another was attributed to ecophysiological and growth differences, or distinct metabolic machinery for the same metabolic pathway^14^. Given this observation and our findings of the ubiquity of viral hosts from Gammaproteobacteria and Campylobacterota, we investigated patterns of relative abundance in the viruses that infect them. We found that Gammaproteobacteria- and Campylobacterota-infecting viruses reflected abundance patterns of the host they infect (**Figure 4**). For example, Campylobacterota-infecting viruses are abundant in vent deposits at Brothers Volcano site NWC-A, however, there is a shift to more abundant Gammaproteobacteria-infecting viruses at Brothers Volcano NWC-B (**Figure 4**), and this is also reflected in microbial abundance of the host taxa (**Figure 4**). We also identified 55 viruses in clusters from geographically separated vents that had a predicted host. Of these, 25 were predicted to infect Campylobacterota or Gammaproteobacteria (**Supplementary Table 4**), further underscoring the potential of these microorganisms to facilitate viral dispersal in hydrothermal vents. In contrast to these taxa, phyla such as Aenigmatarchaeota, Micrarchaeota, an unknown bacterial phylum (EX4484-52), and Iainarchaeota are low abundance microbial community members (less than 1% relative abundance at all sites; Supplementary Table 9). In line with this, the viruses predicted to infect these microorganisms are few in number (29 viruses total), were not present in the nucleotide clusters (inter or intra vent) and were low in relative abundance.

**Figure 4.**
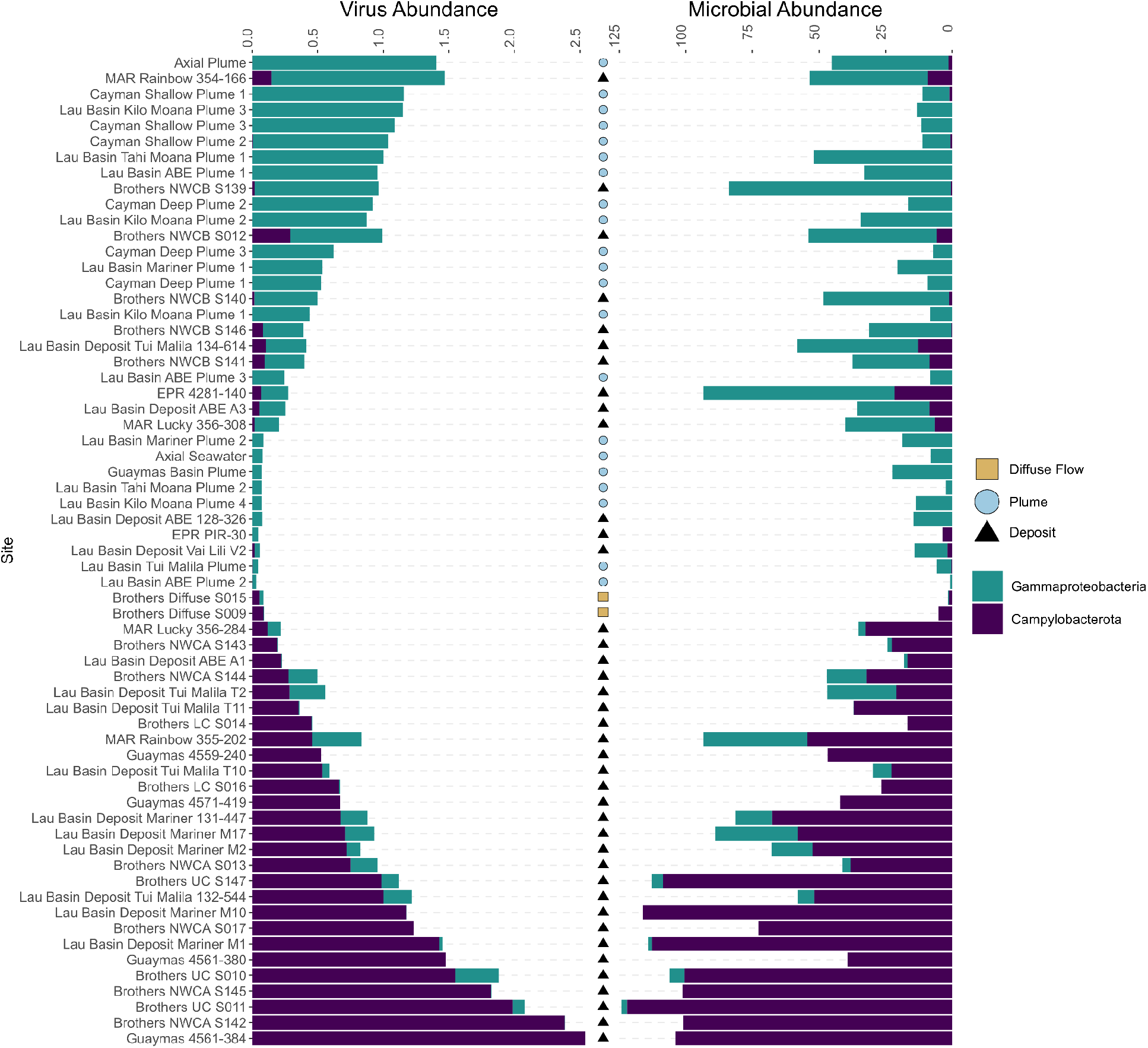
Viral abundance of Gammaproteobacteria- and Campylobacterota-infecting viruses mimics functional redundancy of the hosts. The relative abundance of viruses infecting Gammaproteobacteria and Campylobacterota is shown on the left, while the relative abundance of Gammaproteobacteria and Campylobacterota MAGs is shown on the right. Both abundances are the result of CoverM read mapping normalized by the number of reads in each sample. Sites are shown on the y axis. Colors of the stacked bar plot show the type of host a virus infects (left plot) or the microbial taxa (right plot) and shapes indicate the sample type.

## Discussion

Hydrothermal vent viruses are known to be a key driver shaping microbial communities, yet, they have remained understudied in these ecosystems. By analyzing viruses and microbes recovered from 52 globally distributed hydrothermal vent metagenomes, we show that endemism shapes viral ecology and evolution in deep-sea hydrothermal vents (**Figure 5**). Few prior studies have investigated hydrothermal vent viruses at a global scale^9^, and this is the first comparison of viral communities between hydrothermal vent chimney deposits and plumes.

**Figure 5.**
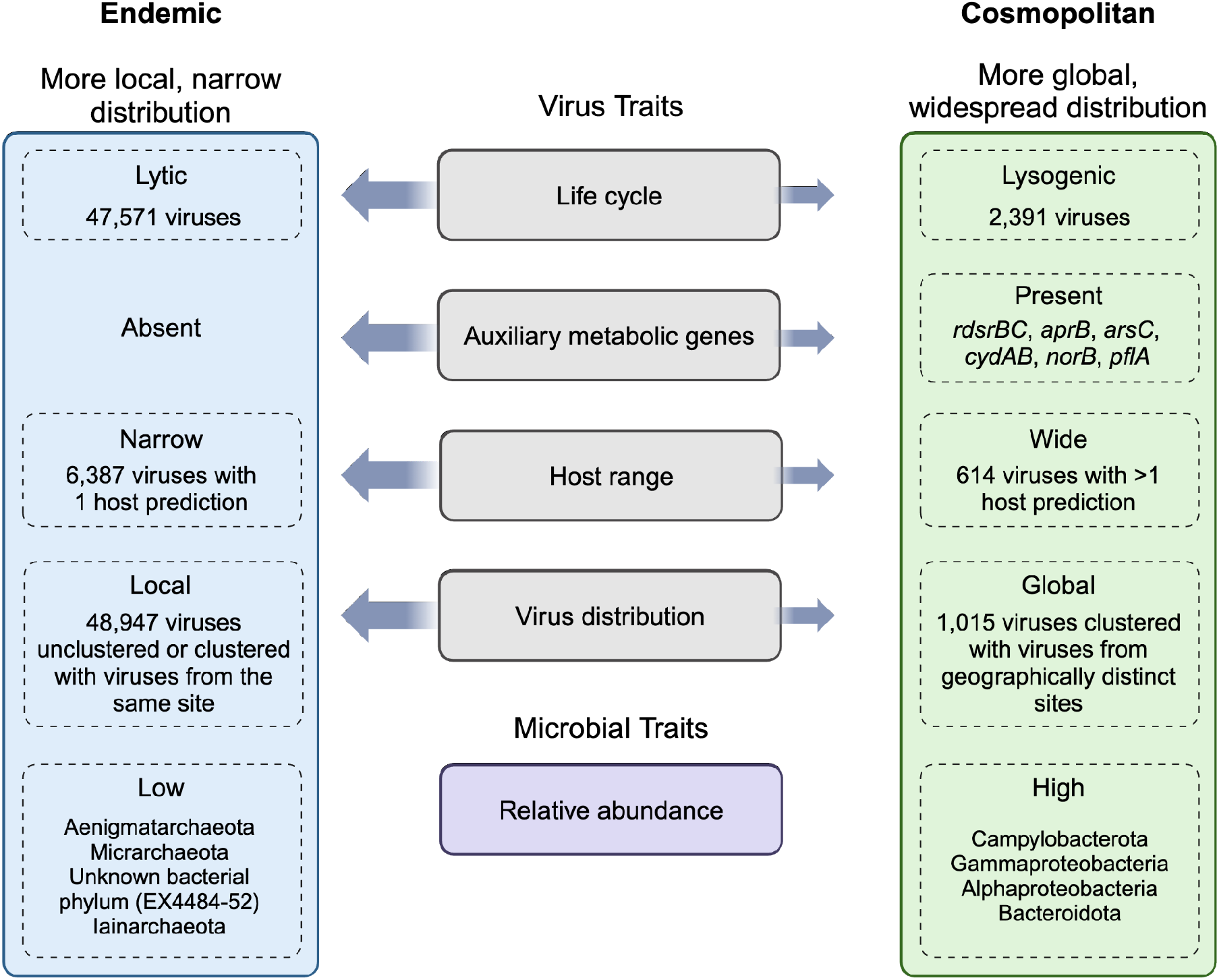
Observed viral and microbial traits suggest endemism shapes the ecology and evolution of viruses in hydrothermal vents. Conceptual diagram showing the viral and microbial traits (center, grey and purple rectangles) observed in this study that contribute to viral biogeography in hydrothermal vents. Arrow width indicates magnitude of support for local versus global viral distribution based on the findings in this study (large width signifies high support, small width indicates low support). The left, blue rectangle shows traits associated with a more local, narrow distribution and the right, green rectangle shows traits associated with a global, widespread distribution. Figure adapted from Chow and Suttle (2015)^12^.

Most viruses identified in this study were characterized as lytic, a viral trait that has been suggested to limit a virus’s distribution compared to lysogeny, where a virus can be dispersed within a host^12^. Though metagenomic-based life cycle predictions are influenced by virus genome completeness, this finding is corroborated by the medium-quality or better viral genomes that we recovered here. The lack of AMGs identified in this study would also limit viral dispersal, as viruses without AMGs would not be as well-equipped to boost energy levels for viral progeny production or reduce the viral latent period^30^. Similarly, microbial host range was narrow for most viruses with a predicted host, which is thought to be limiting in dispersal compared to a wide host range (viruses able to infect >1 genera). Among microbial host traits, Campylobacterota and Gammaproteobacteria MAGs were widespread and abundant, suggesting the viruses infecting these hosts have a greater potential to disperse. Indeed, we found support for this in the relative abundance of viruses infecting these taxa, as well as their presence in nucleotide clusters from geographically separated vents. Conversely, viruses infecting low abundance microorganisms were observed to be rarer and had limited dispersal. Thus, most hydrothermal vent viruses have traits that have been previously hypothesized to promote a narrow, local distribution.

The viral characteristics observed in this study provide a more holistic view of the hydrothermal vents analyzed here, which have previously been characterized for their microbial diversity^13,14,19^. In the hydrothermal plumes, prior studies showed that sulfur compounds dominated as an energy source and plumes consisted of 14 core microbial genera, six of which were within the class Gammaproteobacteria. In deposits, many bacterial and archaeal genera were identified as endemic and Gammaproteobacteria and Campylobacterota were shown to exhibit functional redundancy associated with energy metabolism including those of sulfur, nitrogen, hydrogen, and oxygen. This was hypothesized to be due to differences in the geochemical profiles of different vents, which then selected for ecophysiological and growth differences between taxa^13,14^. These observations are also reflected in the viruses recovered in this study, where Gammaproteobacteria-infecting viruses were abundant in hydrothermal plumes that are typically associated with lower concentrations of hydrothermal compounds such as hydrogen sulfide, sulfur, and hydrogen, while Camplyobacterota-infecting viruses were more abundant in deposits that are associated with higher concentrations of reduced hydrothermal compounds. The viral abundance patterns thus reflected the functional redundancy of Campylobacterota and Gammaproteobacteria in hydrothermal systems. These results underscore the influence of geological context in driving the evolution of microorganisms, and the resulting coevolution of viruses with their microbial hosts. This coevolution likely contributes to high host specificity and the high levels of endemism we observed among viral populations.

The unique viral communities we observed in hydrothermal plumes versus deposits is consistent with previous microbial studies^31^, as well as viral studies that have shown differentiation of viruses between sediments and plumes^9^. While microbial communities in deposits have been found to correlate with geochemistry^13,32,33^, there is evidence that plume microbial communities do not^31^. Instead, microbial communities in plumes of the Lau Basin were shown to be similar, despite differences in their geography, depth, and geochemistry. This aligns with our findings, where the most intra-vent virus similarity was identified in Lau Basin plumes, despite the smaller number of viruses recovered here compared to Brothers Volcano and Lau Basin deposits. Lau Basin plume connectivity was suggested to be promoted by characteristics such as weak stratification and diapycnal mixing over rough topography^31^. Greater sampling resolution is needed within hydrothermal vent fields to dissect the role of local geography in promoting connectivity through hydrothermal plumes. These studies should also be conducted temporally, as hydrothermal systems are dynamic and can change drastically over short time periods based on tectonic activity^34^. Investigations of the same site over time will better elucidate how changes in geology drive coevolution of viruses and microorganisms.

Functionally, viral proteins shared between different vents are predicted to be involved in genome replication, viral structural components, viral infection, lysogeny, and often, are of hypothetical or unknown function. Similar to previous work, our study is hampered by our inability to annotate a large number of viral proteins^35^, where less than half of the proteins shared between different vents were annotated. Poor protein annotation rates inhibit our understanding of the mechanistic underpinnings of connectivity, and thus the unannotated proteins identified here may represent interesting targets for future work to better understand the core proteome of hydrothermal vent viral communities. Of those annotated, few of the proteins shared between geographically separated vents have auxiliary functions, such as those related to microbial dissimilatory metabolism, and these genes were not common in the dataset overall. Thus, although viral genes related to microbial energy metabolism pathways such as sulfur oxidation are known to occur in hydrothermal vents^8^, these genes appear to be rare in these ecosystems.

The high levels of endemism identified in hydrothermal vent viral communities in this study and prior work^9,11^, combined with endemism identified in vent microorganisms and animals^36^, suggests these ecosystems could be especially negatively impacted by future disturbance^37^. In the face of deep-sea mining and anthropogenic climate change, microbial diversity, biomass, and metabolic rates may be severely negatively impacted^38^, and this in turn will be detrimental to viral communities. The large number of unknown viral proteins highlights the vast biological potential we stand to lose as a result. In the future, additional studies are needed to probe the functions of unknown viral proteins, investigate biogeographic patterns on a temporal scale, and obtain a better sampling resolution of hydrothermal vents. This will provide a better understanding of how deep-sea mining and other anthropogenic influences will impact hydrothermal vent communities, and how we can mitigate these disturbances.

## Methods

### Sample collection

Hydrothermal plume samples were collected from the corresponding cruises (Supplementary Table 1): R/V New Horizon in Guaymas Basin, Gulf of California (July 2004)^39– 41^, R/V Atlantis in Mid-Cayman Rise, Caribbean Sea (Jan 2012)^40^, R/V Thomas G Thompson in the Eastern Lau Spreading Center (ELSC) western Pacific Ocean (May-July 2009)^8,40^, and R/V Thomas G Thompson in Axial Seamount, Juan de Fuca Ridge, northeastern Pacific Ocean (Aug 2015)^42^.

Guaymas Basin plume samples were collected by “tow-yo” casts using a CTD rosette in 10 L Niskin bottles^41^. This water was then filtered onto 142 mm 0.2 µm polycarbonate filters by N_2_ gas pressure filtration and preserved in RNAlater^43^. Mid-Cayman plume samples were collected using a Suspended Particle Rosette Sampler (SUPR) by filtering 10-60 L plume water onto 142 mm 0.2 µm SUPOR membranes^40^. These samples were then preserved in RNAlater *in situ*. In Lau Basin, SUPR-collected samples were filtered onto 0.2 and 0.8 μm pore size SUPOR polyethersulfone membranes *in situ* and preserved in RNAlater-flooded vials^31^. In Axial Seamount, plume samples were collected by a Seabird SBE911 CTD and 10 L Niskin bottles^42^. Samples of 3 L were then transferred into cubitainers and filtered through 0.22 μm Sterivex filters.

Hydrothermal deposit samples were collected from the corresponding cruises: R/V Thomas G Thompson in Brothers Volcano, western Pacific Ocean (March 2018)^13^, R/V Roger Revelle (April and May 2015)^13^ and R/V Melville (April 2005)^14^ in the ELSC, western Pacific Ocean, R/V Atlantis in Guaymas Basin, Gulf of California (Nov and Dec 2009)^14,44^, R/V Atlantis in the Mid-Atlantic Ridge, Atlantic Ocean (July 2008)^14^, and R/V Roger Revelle in the East Pacific Rise, Pacific Ocean (March 2004 and December 2006)^14^. Once on ship, deposit samples were subsampled with the outer few millimeters (up to approximately 5 mm) kept separate from the bulk sample. These exterior samples were homogenized and stored at −80°C for subsequent DNA extraction^33^.

### DNA extraction and sequencing

For Guaymas Basin, Mid-Cayman Rise, and Lau Basin plume samples, DNA was extracted from ¼ filters using chemical and physical lysis methods as described in Dick and Tebo (2010)^39^ and Li and Dick (2015)^40^, and sequenced with Illumina HiSeq2000 at the University of Michigan DNA Sequencing Core. Axial Seamount plume samples were extracted using a phenol chloroform extraction and metagenomic libraries were constructed using the Ovation Ultralow Library DR multiplex system^42^. Sequencing was completed using a NextSeq 500 at the W.M. Keck sequencing facility, Marine Biological Laboratory, in Woods Hole, MA.

For Brothers Volcano and ELSC (2015) deposit samples, DNA was extracted from homogenized deposits using the DNeasy PowerSoil kit (Qiagen) and metagenomic libraries were constructed using Nextera DNA Library Prep kits (Illumina), as described in Reysenbach et al., (2020)^13^. Sequencing was completed at the Oregon State University Center for Genome Research and Computing on an Illumina HiSeq 3000. For ELSC (2005), MAR, EPR, and Guaymas Basin, DNA was extracted using the Ultra Clean Soil DNA Isolation Kit (MoBio Laboratories, Carlsbad, CA, USA)^33^. Metagenomic libraries were prepared and sequenced at the Department of Energy, Joint Genome Institute (JGI)^14^.

### Metagenomic assembly and microbial binning

Hydrothermal plume assemblies and microbial MAGs were generated as described in Zhou et al., 2023^19^. Briefly, metagenomic assemblies were constructed from QC-processed reads with MEGAHIT v1.1.2^45^ using the following parameters: -- k-min 45 --k-max 95 --k-step 10. Plume assemblies from Mid-Cayman Rise, Lau Basin Abe, Mariner, and Tahi Moana represent combined plume and background seawater. In other words, for these samples, plume reads were co-assembled with background seawater reads. Microbial MAGs were generated using MetaBAT v0.32.4^46^ using 12 combinations of parameters, followed by DAS Tool v1.0^47^ to generate consensus MAGs. Following MAG refinement and contaminant removal, only MAGs with >50% completeness and <10% contamination were retained, as determined by CheckM v1.0.7^48^.

Hydrothermal deposit assemblies and microbial MAGs were generated as described in Zhou et al., 2022^14^. Briefly, reads from Brothers volcano and ELSC (2015) were quality-filtered using FastQC v0.11.8 (https://www.bioinformatics.babraham.ac.uk/projects/fastqc/) and de novo assembled using metaSPAdes v3.12.0^49^ with the parameters: -k 21,33,55,77,99,127 -m 400 – meta. Reads from ELSC (2005), MAR, EPR, and Guaymas Basin were assembled by the Department of Energy, Joint Genome Institute (JGI) using metaSPAdes v3.11.1^49^ with the following parameters: -k 33,55,77,99,127 –only-assembler –meta. MetaWRAP v1.2.2^50^ was used to generate microbial MAGs with parameters –metabat2 –metabat1 –maxbin2. DAS Tool v.1.0^47^ was then used to generate consensus MAGs.

### Virus identification and binning

VIBRANT v1.2.1^51^ was run with default parameters to identify viruses from the genomic assemblies of the 52 hydrothermal vent samples, resulting in 64,220 viral scaffolds. Viral scaffolds were binned using vRhyme v1.1.0 on each of the 52 hydrothermal vent samples with default parameters and bam files^18^. The sorted bam files used in binning were generated by mapping the fastq reads for a particular sample to the genomic assembly reconstructed for the same sample. Specifically, a custom python script was used to run BWA-MEM v0.7.17^52^ to map reads to assemblies and samtools v1.7^53^ to convert sam files to bam format and then obtain sorted bam files. In total, we reconstructed 38,014 viral genomes. vMAGs were screened for high protein redundancy and binning of lytic and lysogenic viruses, where those with ≥2 redundant proteins and/or ≥2 lysogenic scaffolds were broken back into individual scaffolds and retained in the dataset as unbinned viruses. Finally, vMAGs >10 scaffolds were retained in the dataset as unbinned viruses.

For finalized vMAGs, the vRhyme script link_bin_sequences.py was used with default parameters to generate one scaffold vMAGs, where each scaffold is linked by 1,500 Ns. This is required by some downstream tools that expect viral genomes to be on one scaffold (e.g., CheckV, iPHoP) as described below. Viral genome size was determined using SeqKit v2.6.1 on unbinned viral scaffolds and binned viruses without N-links^54^. To visualize the number of viruses reconstructed from each site, Figure 1 was generated using a custom R script available at: https://github.com/mlangwig/HydrothermalVent_Viruses/tree/main/SitesMap. Viruses were designated as lytic or lysogenic based on VIBRANT, which uses the presence/absence of integrase or the excision of a viral region from metagenomic scaffolds to determine whether a virus is lytic or lysogenic. For vMAGs that had lytic and lysogenic scaffolds binned into one genome, the genome was designated as lytic (https://github.com/mlangwig/HydrothermalVent_Viruses/blob/main/VentVirus_Analysis/VentVirus_Analysis.R).

### Virus taxonomy, marker genes, and quality

Virus taxonomy was determined using geNomad v1.5.1^55^, which utilizes taxonomically informed marker genes to determine the most specific taxon supported by most of the viral genes in the genome. Taxa are defined according to the International Committee on the Taxonomy of Viruses (ICTV)^56^. The end_to_end pipeline was run with default parameters. This also allowed us to determine the number of viral hallmarks encoded by each viral genome. Virus genome quality was determined using the CheckV v0.8.1 end_to_end pipeline with default parameters^57^. For both geNomad and CheckV, vMAGs were input with scaffolds concatenated by 1,500 Ns.

### Virus nucleotide clustering

The average nucleotide identity (ANI) of ≥3kb viruses was calculated with skani v0.2.0^17^ and clustered using the Markov Clustering Algorithm (mcl, release 14-137)^58^. Vskani v0.0.1 (available at https://github.com/cody-mar10/skani-vMAG) was used to run skani and mcl, treating vMAGs as genomes and unbinned viruses as single scaffolds, with the following parameters: vskani skani -c unbinned_PlumeVent_viruses.fna -d fna_vMAGs -x .fasta -m 200 -cm 30 -s 70 -f 50 -ma .7. Option -m signifies the number of marker k-mers used per bases, and was lowered to 200 from the default 1,000 due to the smaller genome size of viruses compared to microorganisms. The compression factor (parameter -cm, equivalent to -c or --slow in skani) was lowered from the default of 125 to 30 to provide more accurate estimates of aligned fractions (AF) for distantly related viral genomes. The screen parameter (-s) removed pairs with less than 70% identity, while the minimum aligned fraction parameter (-f) kept ANI values where one genome had an aligned fraction greater than or equal to 50%. Finally, -ma signifies minimum ANI, which was lowered to 70% from the default 95% to capture a broader range of viral relatedness (the the family and genus level). Mcl clustering was completed using default parameters.

The resulting table of skani-produced ANI and AF values was manipulated using a bash script, where ANI was normalized by the lowest AF (ANI*AF/100^2^), filtered for ≥70% ANI (now corrected for AF), and formatted as input for the mcxload function of mcl (https://github.com/mlangwig/HydrothermalVent_Viruses/blob/main/ANI_clustering/ANI_clust.R). Mcl clustering was then run with default parameters. The resulting cluster file was then input into R to calculate the average ANI per cluster, map metadata to the clusters, and determine which clusters contain viruses from geographically distinct sites (https://github.com/mlangwig/HydrothermalVent_Viruses/blob/main/ANI_clustering/ANI_clust.R).

To identify the regions of overlap between low-quality viruses, BLASTN v2.14.1^59^ was run per cluster on all viruses within a cluster. First, a blast database was made for each cluster with the following command: makeblastdb -in -out -dbtype nucl. Then, blastn was run on each cluster with the following command: blastn -query -db -out -outfmt “6 qseqid sseqid evalue bitscore length pident qstart qend sstart send” -max_target_seqs 2 -max_hsps 1. The aligned coordinates were obtained from these output files. Finally, bedtools^60^ was used to extract these coordinates from the amino acid-format virus genome files with the following command: bedtools intersect -a bed_GeoDistinct_VirusClusts.tsv -b bed_PlumeVent_viruses.tsv -wa -wb > result.txt. The -a bed file contains the coordinates from the blastn results and the -b bed file contains the coordinates of all the ORFs in all the viral genomes. This allowed us to obtain the gene annotations of the regions of the viral genomes that had identical nucleotides.

### Virus protein clustering

Mmseqs2 (v15.6f452)^61^ was used to cluster viral proteins. First, the createdb option was used to create an mmseqs database of the 595,416 hydrothermal vent virus proteins. Next, the cluster option was used with the following parameters: -cov-mode 0 --min-seq-id 0.75. These options signify that the alignment covers at least 80% of the query and of the target, and that the minimum sequence identity is 75%. The default clustering algorithm, greedy set cover algorithm, was used. This algorithm iteratively selects the node with the most connections and all its connected nodes to form a cluster, and repeats this process until all nodes are within a cluster. Next, the createtsv option was used to generate a tsv file of the cluster output. This file was parsed and analyzed in R (https://github.com/mlangwig/HydrothermalVent_Viruses/blob/main/ANI_clustering/Protein_clustering.R).

### Protein annotations

Viral proteins were annotated using VIBRANT^51^, which employs hmmsearch to annotate viral proteins with KEGG, VOG, and pfam HMM databases. To determine the best supported VIBRANT hits from the three databases, the annotation with the highest bit score was chosen, followed by the lowest e-value, and finally the highest viral score. Viruses were also annotated using DRAMv v1.4.6^62^ to identify potential AMGs. To run DRAMv, VIBRANT-identified viruses were input into Virsorter2 v2.2.4^63^ to obtain the input file needed for the DRAMv software. Because Virsorter2 was used for downstream analyses and not viral discovery, we used the following parameters: virsorter run --keep-original-seq --prep-for-dramv --include-groups dsDNAphage,NCLDV,RNA,ssDNA,lavidaviridae --provirus-off --viral-gene-enrich-off --min-score 0.0. DRAMv annotate was run with the default parameters. DRAMv distill was run with default parameters to obtain annotations that are supported as AMGs.

### Read mapping

All 49,962 viral genomes were mapped to all 163 paired-end fastq reads using CoverM v0.6.1 (https://github.com/wwood/CoverM) with the options --methods count relative_abundance --min-covered-fraction 0. For read mapping used to determine connectivity between vents, the output was filtered in R to only retain viruses where reads mapped to ≥70% of the viral genome (https://github.com/mlangwig/HydrothermalVent_Viruses/blob/main/Read_Mapping/Coverm_circos.R). To obtain normalized relative abundance, the number of reads mapped to a virus from each sample was divided by the number of reads in that respective sample. This methodology was repeated with the microbial MAGs to obtain their normalized relative abundances.

### Host prediction

iPHoP v1.3.3 was used to identify virus-host links between hydrothermal vent viruses and a custom database of 3,872 MAGs reconstructed from the same sites^14,19,64^. Before building the custom database, BLASTN^59^ was used to search all viruses against all microbial MAGs with the following command: nohup blastn -query -db -out -outfmt “6 qseqid length qlen slen pident bitscore stitle”. Hits with 100% identity and 100% coverage were considered viral contamination and were removed from microbial MAGs. Following this step, the custom MAG database was created using the add_to_db option and iPHoP was run using the following parameters: iphop predict --db_dir --no_qc. The input file included unbinned viral scaffolds and vMAGs concatenated into one scaffold using 1,500 Ns to enable one prediction per vMAG.

## Data Availability

Genomic assemblies and microbial metagenome-assembled genomes were previously published through NCBI BioProject IDs PRJNA488180 and PRJNA821212. Viral genomes are available at https://figshare.com/articles/dataset/Hydrothermal_Vent_Viruses/25968037. Scripts used in this work are available at https://github.com/mlangwig/HydrothermalVent_Viruses.

## Author contributions

MVL, ALR, and KA conceptualized the project. KA supervised the project. KA and ALR obtained and sequenced the hydrothermal samples. ZZ performed metagenomic assembly and binning. MVL identified viruses from the assemblies, performed viral binning, and all downstream analyses. MVL and FK analyzed viral AMGs. CM developed software for analyses. MVL conducted data validation, curation, analysis, created visualizations, and administered the project. MVL and KA wrote the manuscript. All authors reviewed the results, edited, and approved the manuscript.

## Acknowledgements

This research was supported by the National Science Foundation under grant numbers DBI2047598 (to KA), OCE2049478 (to SBJ, KA) and OCE-0728391, OCE-0937404, OCE-1558795 (to ALR). CM was funded by a National Science Foundation Graduate Research Fellowship. We thank the crew of the R/V *Roger Revelle*, R/V *Atlantis*, R/V *Thomas G. Thompson*, HOV *Alvin*, and the ROV *Jason* for assistance in collecting these samples. Thank you to Spencer R. Keyser for your help with data wrangling in R.

