## Supplementary Text for "Endemism shapes viral ecology and evolution in globally distributed hydrothermal vent ecosystems"

### Geographically distant hydrothermal vents share viruses

Viruses in the 65 geographically distinct clusters are predominantly predicted to be low-quality. Because these viruses have low estimates of genome completeness, we wanted to understand which regions of the viruses overlap and confirm that they are biologically meaningful. In a cluster of three low-quality, putatively lytic viruses from the Lau Basin and Brothers Volcano, the viral genomes overlap up to 21 kb at 93.2% identity, and this spans a minor tail protein and major capsid protein (**Supplementary Table 5**). In more distant sites, a viral pair reconstructed from the plumes of Axial Seamount and Guaymas Basin, respectively, shared 6.1 kb at 99.7% identity, and this region included six VOGs, one of which is a tail assembly chaperone. In another cluster, two short, low-quality viruses from the Mid-Atlantic Ridge and Brothers Volcano share 96.6% identity across a 2.9 kb region, which includes two baseplate proteins. Thus, even among low-quality viruses, we find support for shared viral genomic regions between geographically separated vents.

### Viral biogeography is closely tied to the geographic distribution and abundance of their hosts

In plume samples, predictions within phylum Pseudomonadota are largely to Gammaproteobacteria, Alphaproteobacteria, and Bacteroidota. Predicted Gammaproteobacteria hosts span a range of genera, including *Colwellia*, *Acinetobacter*, *Thioglobus*, and *Alteromonas*, all of which are known to occur in hydrothermal plumes and seawater[1–4]. Predicted Alphaproteobacteria hosts include families Pelagibacteraceae (SAR11) and Sphingomonadaceae, supporting previous findings of the dominance of these organisms in seawater and hydrothermal plumes[5]. Similarly, most Bacteroidota host predictions are to the family Flavobacteriaceae, which has been identified in hydrothermal plumes with abundant extracellular peptidase genes, used to acquire carbon and nitrogen from the environment[6,7]. Compared to 1,131 predictions to bacterial hosts in the plume, there are 87 predictions to archaeal hosts and more than half of these are predictions to orders Nitrososphaerales, Pacearchaeales, and Poseidoniales. These orders have previously been identified as dominant archaea in hydrothermal plumes and deep seawater[8–10]. Pacearchaea in Guaymas Basin hydrothermal plumes have been characterized as having high connectivity with other microbial groups based on metabolic networks, and are predicted to receive the greatest benefits from community interactions, including cellobiose, oxygen, carbon dioxide, and sulfide[10]. Microorganisms in the order Nitrososphaerales are known to widely encode *amoA* for ammonia oxidation (Thaumarchaea, formerly Marine Group I)[11] and Poseidoniales (formerly Marine Group II) that reside in the deep ocean are predicted to reduce nitrate, though are still largely understudied[8].

Viruses are predicted to infect a greater diversity and larger number of microbial phyla in hydrothermal deposits compared to plumes (6,591 host predictions in deposits, Supplementary Table 2). Most host predictions within phylum Pseudomonadota are to Campylobacterota (1,679), Gammaproteobacteria (685), and Alphaproteobacteria (675). More than half of the predictions to Campylobacterota are within the genera *Sulfurovum*, *Sulfurimonas*, and a genus with no cultured representatives, *UBA1140*. Cultured *Sulfurovum* and *Sulfurimonas* isolates from hydrothermal vents are known to be chemolithoautotrophic with the ability to oxidize sulfur and hydrogen and reduce sulfur, nitrate, and thiosulfate[12–14]. Among Gammaproteobacteria, some viruses are predicted to infect host genera that were also found in the plume, including *Colwellia*, *Acinetobacter*, and *Alterimonas*, however, the number of predictions to these microorganisms were much smaller. In deposits, predicted Gammaproteobacteria hosts largely include unknown genera, *Thiolapillus*, *Cocleimonas*, *Thiogranum*, and *Thiomicrorhabdus*. All of these bacterial genera have cultured representatives isolated from hydrothermal vents or deep sea sediments and are capable of chemolithoautotrophic sulfur oxidation[15–18]. Finally, Alphaproteobacteria host predictions are also largely to unknown or undescribed genera (*UBA5972* and *UBA3077*), as well as *Profundibacter* and *Thermopetrobacter*. *Profundibacter* was originally isolated from Loki's Castle vent field and is known to be piezophilic and anaerobic[19], while *Thermopetrobacter* was cultured from the Eastern Lau Spreading Center and is an aerobe capable of chemoautotrophic growth on hydrogen[20].

Of the 6,591 viral host predictions in deposit samples, 875 are attributed to Archaea, largely to phyla Thermoproteota (348), Methanobacteriota_B (195), Halobacteriota (111), and Thermoplasmatota (77). Thermoproteota are known to be common in deep-sea hydrothermal vent archaeal communities, though they are underrepresented in genomic databases[21]. In our dataset, nearly half of the Thermoproteota predictions are within families Acidilobaceae and Desulfurococcaceae. Acidilobaceae have recently been amended to include eight genera and range from acidophilic to neutrophilic thermo- or hyperthermophiles that use carbohydrates or protein-rich carbon for growth[21]. The Desulfurococcaceae family are also hyperthermophiles, capable of growing heterotrophically by sulfur respiration of organic compounds or chemolithoautotrophic sulfur reduction with hydrogen as the electron donor[22]. Nearly all virus-host predictions to phylum Methanobacteriota_B are within the *Thermococcus* genus (173/193). These archaea are ubiquitous in hydrothermal vents, where they are known to be sulfur-reducing hyperthermophiles and have the ability to use mixed heterotrophic and carboxydotrophic metabolism[23]. Among Halobacteriota, many host predictions are to the closely related *Archaeoglobus_B*, *Archaeoglobus_C*, and *Geoglobus* genera. Cultured isolates of *Archaeoglobus_B* and *Archaeoglobus_C* are strict anaerobic hyperthermophiles that use sulfur compounds as terminal electron acceptors, but are distinguished by an incomplete Wood-Ljungdhal pathway and inability to reduce sulfate, respectively[24]. The *Geoglobus* genus is represented by two cultured isolates, isolated from the Guaymas Basin and the Mid-Atlantic Ridge, which are hyperthermophilic, anaerobic, and Fe(III)-reducing archaea[25]. Thermoplasmatota host predictions are mostly to undescribed genera, with the exception of *Aciduliprofundum*, whose sole cultured representative was isolated from hydrothermal vents and is an anaerobic heterotrophic sulfur- and iron-reducing thermoacidophile[26].
