## Supplementary Figures for "Endemism shapes viral ecology and evolution in globally distributed hydrothermal vent ecosystems"

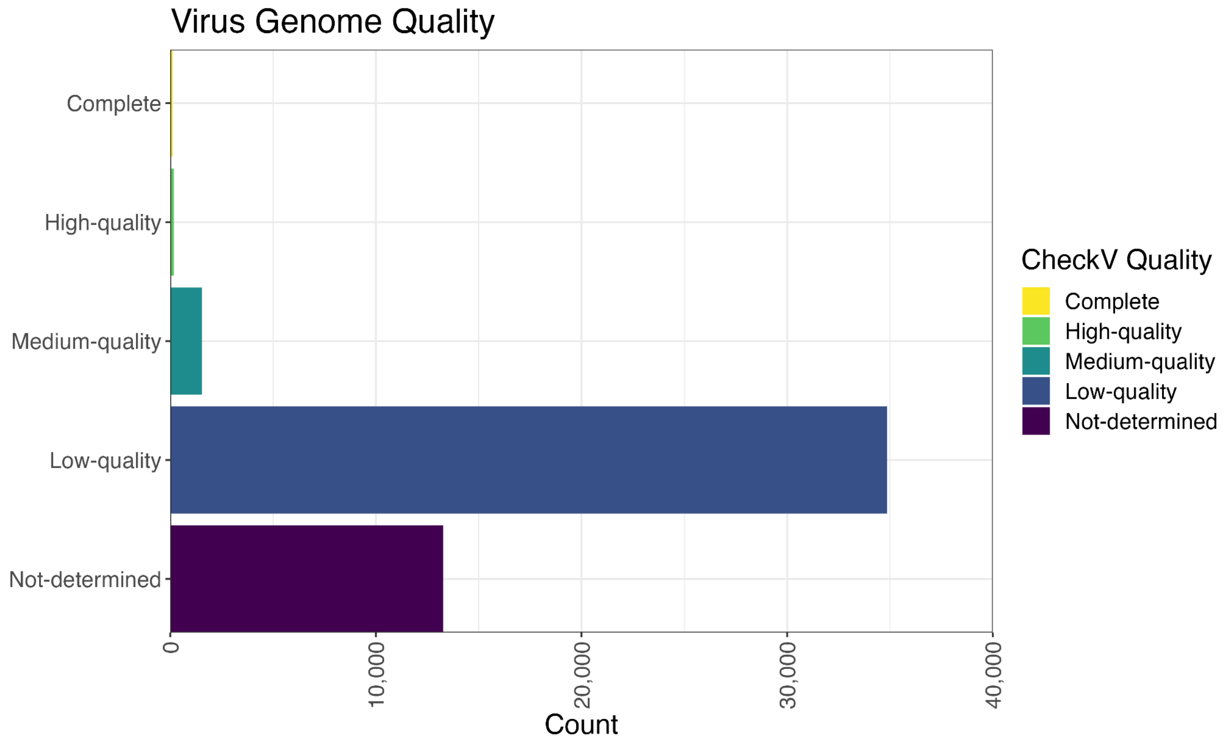


**Supplementary Figure 1**. Genome quality estimates of the 49,962 hydrothermal vent viruses. Figure was generated in R. (https://github.com/mlangwig/HydrothermalVent_Viruses/blob/main/VentVirus_Analysis/VentVirus_Analysis2.R).

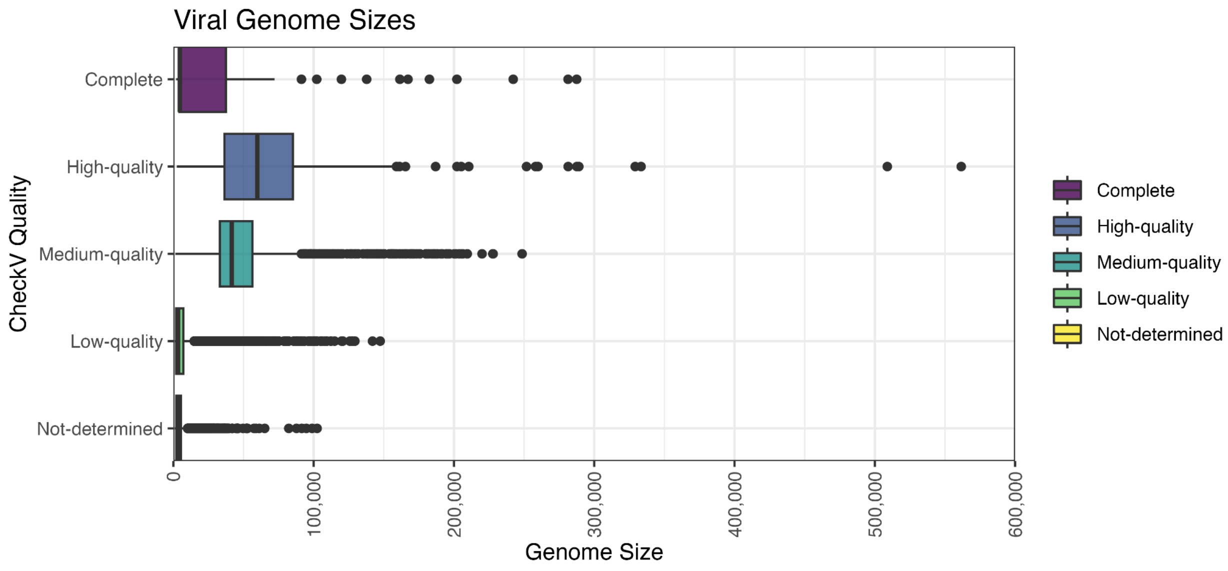


**Supplementary Figure 2.** Virus genome size distribution for each quality estimate category. Viral genome size was determined using seqkit. Figure generated in R (https://github.com/mlangwig/HydrothermalVent_Viruses/blob/main/VentVirus_Analysis/VentVirus_Analysis2.R).

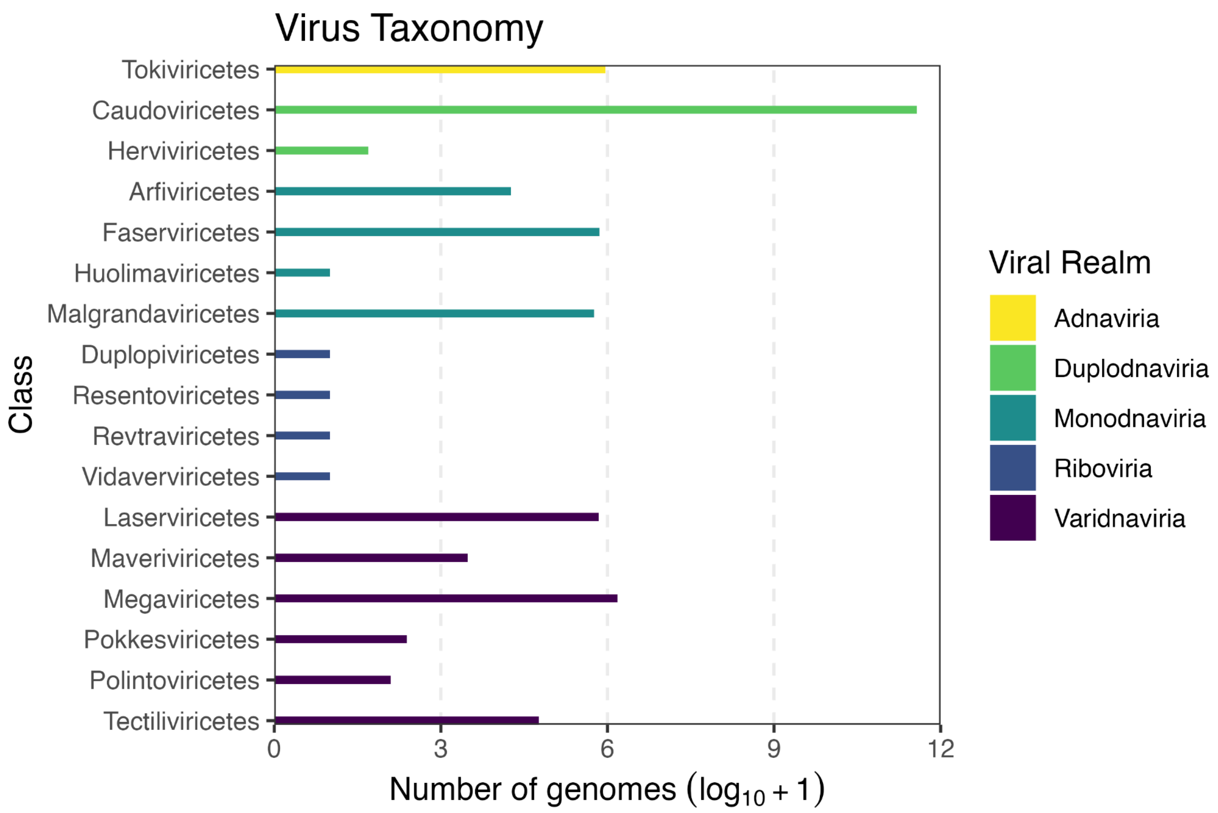


**Supplementary Figure 3.** Virus taxonomy of 40,236 vent viruses. The x axis shows the log+1-transformed number of viral genomes in each class. Bars are colored according to the realm of each class. Viruses that were assigned to an unknown realm and/or class are not included (392 viral genomes). Figure generated in R (https://github.com/mlangwig/HydrothermalVent_Viruses/blob/main/VentVirus_Analysis/VentVirus_Analysis2.R)

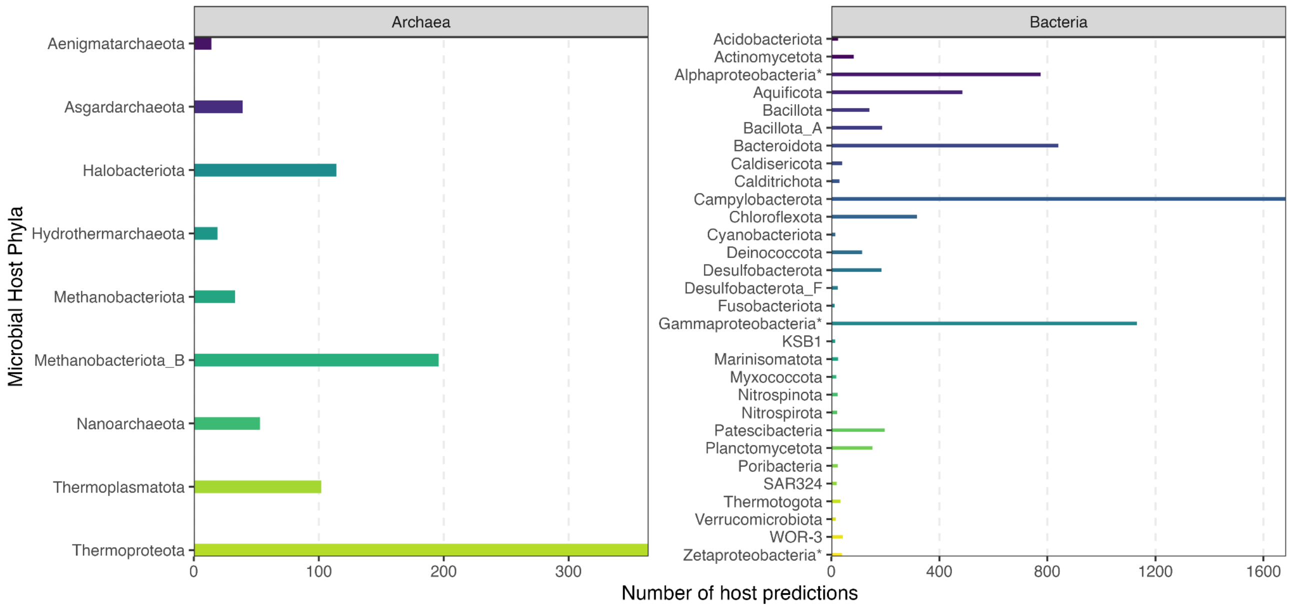


**Supplementary Figure 4.** Microbial host predictions for 7,001 hydrothermal vent viruses. Microbial phyla are shown on the y axis and the total number of hosts are shown on the x axis. Plots are faceted according to the microbial domains archaea and bacteria. Host predictions were made using iPHoP.  Figure generated in R (https://github.com/mlangwig/HydrothermalVent_Viruses/blob/main/VentVirus_Analysis/VentVirus_Analysis2.R)
